# Evolutionary Models of Amino Acid Substitutions Based on the Tertiary Structure of their Neighborhoods

**DOI:** 10.1101/468173

**Authors:** Elias Primetis, Spyridon Chavlis, Pavlos Pavlidis

## Abstract

Intra-protein residual vicinities depend on the involved amino acids. Energetically favorable vicinities (or interactions) have been preserved during evolution, while unfavorable vicinities have been eliminated. We describe, statistically, the interactions between amino acids using resolved protein structures. Based on the frequency of amino acid interactions, we have devised an amino acid substitution model that implements the following idea: amino acids that have similar neighbors in the protein tertiary structure can replace each other, while substitution is more difficult between amino acids that prefer different spatial neighbors. Using known tertiary structures for *α*-helical membrane (HM) proteins, we build evolutionary substitution matrices. We constructed maximum likelihood phylogenies using our amino acid substitution matrices and compared them to widely-used methods. Our results suggest that amino acid substitutions are associated with the spatial neighborhoods of amino acid residuals, providing, therefore, insights into the amino acid substitution process.

## 1 Introduction

Structure and functionality of a protein are closely related (Worth et al., 2009). Its amino acid sequence as well as cofactors, ligands and other parts of the same or other proteins form a complex network of interactions that is the basis of the unique physicochemical properties of each protein related to its function (Worth et al., 2009). During the last decade, a multitude of tertiary protein structures have been determined by using techniques such as crystallography, NMR, electron microscopy and hybrid methods (Egli, 2010). Consequently, the number of the available tertiary structures in the RCSB-PDB database (Bernstein et al., 1977) amino acid is rapidly increasing.

Computational methods facilitate structural, functional and evolutionary characterization of the proteins Nath Jha et al. (2011). Protein function is tightly linked to its tertiary structure, and consequently to the vicinities, or interactions, amino acid residues have formed. For example, globular and membrane proteins are characterized by totally different physicochemical environments. Membrane proteins show a limited interaction with water molecules, and they are able to interact with the lipid bilayer. Thus, their trans-membrane region adopts a single type of secondary structure – either a helix or a beta-sheet. Largely, the secondary structure is defined by non-neighboring (on the sequence level) amino acid residue interactions (Nath Jha et al., 2011).

Recently, we developed PrInS (freely available from http://pop-gen.eu/wordpress/software/prins-protein-residues-interaction-statistics (Protein Interaction Statistics; Pavlidis et al. unpub-lised) an open-source software to score proteins based on the frequency of their residue interactions. PrInS uses protein structures stored in Protein Data Bank (PDB) to construct a statistical model of intra-protein amino acid residue interactions for a certain class of proteins (e.g. membrane proteins). PrInS scores every amino acid *a* proportionally to the number of ‘unexpected residues’ that interact with it. The term ‘unexpected residues’ means residues that they are rarely found to be in the vicinity of *a* if we consider all protein structures of a given dataset. Therefore, PrInS is able to pinpoint residues characterized by a large number of ‘less frequent’ interactions, and therefore they may represent functional areas of the proteins or targets of natural selection based on the following assumption: even though a large number of unlikely interactions characterizes these amino acids, nature has preserved them. Here, we use PrInS to describe statistically the intra-protein amino acid interactions and consequently to construct an amino acid substitution matrix endowed with the principle that amino acids with similar (residual) neighborhoods can substitute each other during evolution.

Protein evolution comprises two major principles. The first principle suggests that protein structure is more conserved than the sequence (Siltberg-Liberles et al., 2011). The second principle suggests that the physicochemical properties of amino acids constrain the structure, the function and the evolution of proteins (Hatton and Warr, 2015). Protein evolutionary rate is strongly correlated with fractional residue burial (Siltberg-Liberles et al., 2011). This is due to the fact that the core of a protein is mostly formed by buried residues, which often play a crucial role in the stability of the folded structure (Franzosa and Xia, 2008). The three-dimensional structure of the protein determines its evolutionary rate, since most mutations in the core of a protein tend to destabilize the protein.

Within protein families the backbone changes are infrequent, thus, preserving the folding properties over relatively long evolutionary distances, while substitutions are found often at the side chains. For example, for proteins with binding function, the binding interface is under functional constraint and may evolve the slowest, with differences in rate between affinity-determining and specificity-determining residues (Siltberg-Liberles et al., 2011).

In addition, the secondary structural elements of a protein evolve at different rates. Beta sheets evolve more slowly than helical regions and random coils evolve the fastest (Siltberg-Liberles et al., 2011). Secondary structure changes may eventually occur due to varying helix/sheet propensity. Some of these changes in secondary structural composition may be evolutionary neutral, whereas some structural transition may involve negative or positive selection. In the latter case, a new mutationally accessible fold may enable the development of a new favorable function that was not possible within the previous fold (Siltberg-Liberles et al., 2011).

In the context of folding, the thermodynamic stability of the proteins with a stable unique tertiary structure is important. Thermodynamic stability is maintained throughout evolution despite the destabilizing effect of the non-synonymous mutations, which are often removed from the populations as a result of negative selection. The protein structure is important because it acts as a scaffold for properly orientating functional residues, such as a binding interface and a catalytic residue. As a result, the selective pressure for particular sequences (and not structures) over longer evolutionary periods is decreased, generating a neutral network of sequences interconnected via mutational changes (Siltberg-Liberles et al., 2011).

Choi and Kim (2006), based on the most common structural ancestor (CSA), showed that not all present-day proteins evolved from one single set of proteins in the last common ancestral organism, but new common ancestral proteins were born at different evolutionary times. These proteins are not traceable to one or two ancestral proteins, but they follow the rules of the ”multiple birth model” for the evolution of protein sequence families (Choi and Kim, 2006).

In this study we used the scoring matrices that are obtained from the PrInS algorithm to examine the evolution of proteins and we focused on the evolution of *α*-helical membrane proteins. The main hypothesis is that protein evolution is related to the three dimensional neighborhoods of amino acids. Specifically, amino acids that have similar amino acid neighbourhoods can substitute each other during evolution.

## 2 Methods

### 2.1 Dataset retrieval and name conversion

In eukaryotes, *α*-helical proteins exist mostly in the plasma membrane or sometimes in the outer cell membrane. In prokaryotes, they are present in their inner membranes. We used 82 *α*-helical membrane proteins, which were also scrutinized previously by Nath Jha et al. (2011) to describe the statistical properties of amino acid interactions within *α*-helical membrane proteins. First, for each of the protein in the dataset, we downloaded multiple sequence alignments of homologous protein sequences from the UCSC Genome Browser (Kent et al., 2002) for 16 primate and 3 non-primate mammalian species. Second, three-dimensional structures were downloaded from the Protein Data Bank (PDB) database.

### 2.2 The PrInS software

We applied PrInS on the three dimensional structures downloaded from PDB. PrInS constructs a scoring matrix *M* as follows: If the tertiary structure of a protein is represented by *P* and the total number of amino acid residues is *l*, then residues 1 ≤ *k,m* ≤ *l* interact if and only if:

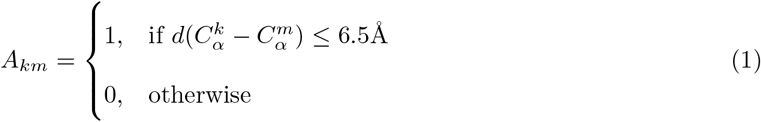

where 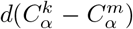 denotes the distance between the *C_α_* atoms of the *k* and *m* amino acids. In other words, the *k^th^* and the *m^th^* residues of a protein interact if and only if the Euclidean distance between their *C_α_* atoms is less than 6.5A and they are not located on adjacent positions on the amino acid chain.

#### 2.2.1 Defining the environment of each amino acid

According to Nath Jha et al. (2011), amino acids of a HM protein can be classified in three environments: The first environment comprises amino acids at different distances from the lipid bilayer. The second environment classifies the helices (and thus, their amino acids) on the basis of their inter-helical interactions inside the membrane. Finally, the third environment classifies amino acid residues based on the number of interactions they make. In our study we have used the third type of environments (residue-contact-based interaction) based on the results from Nath Jha et al. (2011), who demonstrated that the residue-contact-based environment description of residue interaction is more accurate for the *α*-helical proteins they studied. Thus, every amino acid residue *P_j_* in protein *P* can be assigned to the pair (*A, K*), where *A* indexes the amino acid type of the *k^th^* residue (e.g. Alanine) and *K* represent the environment of the *k^th^* residue. Based on the results of Nath Jha et al. (2011), we used the number of non-covalent contacts each amino acid makes to define its environment. Thus, environment I comprises all amino acids with 1-5 contacts, environment II, amino acids with exactly 6 contacts and environment III amino acids with more than 6 contacts.

Initially, PrInS was used to construct a matrix M, a 60 × 60 matrix (or equivalently nine 20 × 20 scoring matrices), that scores amino acid interactions in all environment pairs. For example, *M_ij_,i,j* ≤ 20 describe score interactions between amino acids of the environment I. In *M* the pairs of amino acids with the lower scores are those that interact frequently. In general, the interaction between amino acid A(1 ≤ *A* ≤ 20) and amino acid B (1 ≤ *B* ≤ 20) that belong to the environments *K* (1 ≤ *K* ≤ 3) and *Q* (1 ≤ *Q* ≤ 3), respectively, is given by:

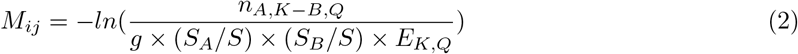

The coordinates *i* and *j* are given by *i* = 20(*K* − 1) + *A* and *j* = 20(*Q* − 1) + *B*, respectively. The parameter *g* = 2 if the contacting pair comprises different amino acids in the same environment (A = *B,K* = *Q*), and *g* =1, otherwise. *S_A_* and *S_B_* are the total number of amino acids *A* and *B* in the dataset, respectively. *S* is the total number of amino acids in the dataset and *E_K,Q_* equals to the number of interactions between environment *K* and *Q*. Finally, *n_A,K−B,Q_* is the number of interactions between amino acids *A* in environment *K* and amino acid *B* in environment *Q*. It is evident that assumes large (positive) values when the observed number of interactions *n_A,K−B,Q_* between *A* and *B*, is much lower than the interactions expected based on the total frequency of the amino acids *A* and *B* in the dataset and the number of interactions between the *K* and *Q* environments. In other words the value of *M_ij_* is large for rare interactions.

#### 2.2.2 Distance matrix

The *M* scoring matrix was split in nine 20 × 20 matrices and distance metrics (Euclidean, Manhattan and Pearson Squared) between the elements of each matrix (amino acids) were calculated as follows: Let *G_d_* represent the average 20 × 20 distance matrix for the distance metrics *d* (*d* ∈ {Euclidean, Manhattan, Pearson Squared}). We name these matrices neighborhood Euclidean-distance based (*nEd*), *nMd* and *nPd*, respectively, for Euclidean, Manhattan and Pearson Squared. For comparison purposes, we created an additional 20 × 20 distance matrix based on the Hamming distance between a pair of codons showing the minimum number of nucleotide changes required to transform one amino acid to another. Smaller distance values indicate higher similarity between the respective amino acids. Distance matrices have been visualized as heatmaps **(Supplementary Figures 8a, 8b, 8c and 8d)**. Two amino acids that have similar neighborhoods, *i.e*. similar amino acid neighbors in the tertiary structure will, thus, have a small distance (high similarity) between them. Consequently, according to our hypothesis, they will substitute each other during evolution at a higher rate.

#### 2.2.3 Substitution rate matrix

Each of the four distance matrices were transformed into rate matrices *D_r_* by using a similar procedure as in Dayhoff et al. (1978). In particular, the following procedure was followed:

1. Find the maximum value of the distance matrix *D_g_*, *D_max_*.
2. Subtract each matrix element from the *D_max_* and divide by *D_max_*. This will transform *D_g_* to an amino acid ”similarity” symmetric matrix, where 1 denotes maximum ‘similarity’ between two amino acids.
3. Similar to the substitution matrices BLOSUM62 and PAM120, one element of the matrix is defined as a reference. All other elements are scaled proportionally to the reference.
4. The diagonal of the matrix is defined as the negative sum of all other elements of the respective row.

If an amino acid pair is characterized by a large (relative) substitution rate, then they can substitute each other during evolution at a high rate. The substitution rate matrix that is derived with the aforementioned algorithm is symmetric.

#### 2.2.4 Evaluation of PrInS ability to predict amino acid substitutions

For the whole set of multiple alignments, a substitution 20 × 20 matrix *S* was created as follows: Let *B* the set of sites from all sequence alignments that contain only two amino acids. Let |*B*| the number of such sites. Then, if *B_i_*, 0 < *i* < |*B*| is such a site, that contains only amino acids *a* and *b*, *B*[*a*][*b*] is incremented by one. In other words, a cell in the *S* matrix that corresponds to amino acids *a* and *b*, counts how many times we observed a site from the multiple sequence alignments with the amino acids *a* and *b*. The assumption here is that during evolution there was at least one substitutions between *a* and *b* for the specific site of the alignment that contains these two amino acids. Also, the more sites with *a* and *b*, the more frequent the substitution between them.

To evaluate the relation of our neighborhood-based substitution distance matrix *nEd* and the multiple sequence alignments, we calculated the Pearson correlation coefficients between each amino acid in *S* and *nEd*. Negative values of Pearson correlation coefficient values suggest that the amino acid substitutions in the alignments are concordant with the neighborhood-based substitution rates distances (smaller distances in nEd suggest greater neighborhood similarity).

Moreover, we manually analyzed the amino acid pairs with the lowest distance values in the Euclidean, Manhattan and Pearson distance matrices with the tool ‘Common Substitution Tool’ in the ‘Amino Acid Explorer’ (https://www.ncbi.nlm.nih.gov/Class/Structure/aa/aa_explorer.cgi) of NCBI (Bulka et al., 2006). Common Substitution Tool sorts an amino acid list from the most common to the least common substitutions, given an amino acid as input, based on the BLOSUM62 substitution matrix (Henikoff and Henikoff, 1992).

### 2.3 Overall and Site Likelihoods

Likelihood is a function of the parameters of a statistical model for given data. Since substitution rate matrices are part of the model, we compared the per site and the overall likelihood values obtained by our substitution matrices and those obtained by using BLOSUM62 and PAM120. Both the phylogeny and the equilibrium frequencies of the amino acids were considered known in this analysis. Thus, by keeping the remaining parameters the same for both models, we aimed to assess which rate matrix can better explain the observed multiple alignments. As overall likelihood, we defined the total likelihood of the alignment, while a site likelihood is the likelihood of a single position in the amino acid alignment. The phylogenetic tree was retrieved from the UCSC Genome Browser (Kent et al., 2002), and the equilibrium frequencies of amino acids were obtained from the equilibrium frequencies in the BLOSUM62 model. We used the same phylogenetic tree and equilibrium frequencies for all comparisons.

## 3 Results

### 3.1 Evaluation of PrInS Ability to Predict Amino Acid Substitutions

The substitution matrix counts the amino acid substitutions that occurred in the 223 downloaded multiple alignments using the two most frequent amino acid residues at each alignment site. For each amino acid in this matrix, we evaluated its correlation (Pearson correlation coefficient) with the same amino acid in the distance matrices. Thus, if, for a given amino acid, both matrices suggest similar substitution preferences, then neighborhood-based substitution models are concordant with the observed substitutions. Since we used distance matrix in this analysis, negative correlation coefficient indicate concordance. Results suggest that there is concordance between the substitution matrix and our neighborhood-based distance matrices for most of the amino acids. Figure 1 shows the Pearson correlation coefficients for amino acids using the substitution matrix and the *nEd*. The correlation plots between the substitution matrix and other distances are illustrated in the **Supplementary Figures 9a–9c**.

**Figure 1:**
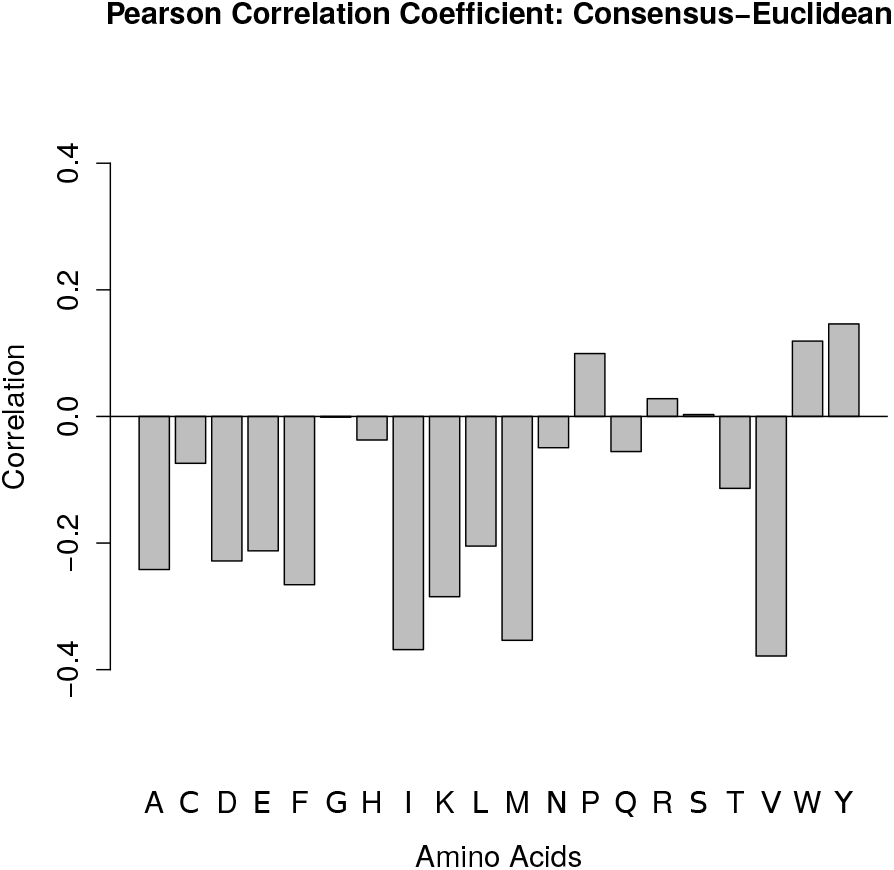
Pearson Correlation Coefficient between the substitution and *nEd*. Concordance is indicated with negative values, while discordance with positive values.

As shown in Figure 1, most of the amino acids are concordant when we compare the Euclidean distance matrix and the substitution matrix.

In the **Supplementary Figure 9b** (substitution vs Manhattan distance), the discordant amino acids were the five out of six amino acids of Figure 1. Similar results are shown in **Supplementary Figure 9c**, where the substitution matrix is compared against the squared Pearson distance matrix. Finally, there are no discordant amino acids in the correlation between the Consensus matrix and the Genetic Code matrix (**Supplementary Figure 9a**).

Furthermore, the amino acid pairs with the lowest values in the distance matrices are at the top of the amino acid list that the Common Substitution Tool of NCBI Amino Acid Explorer reports. This means that amino acids with similar three dimensional neighborhoods tend to substitute each other. For example, in Euclidean distance matrix, the Leukine-Isoleukine pair has the lowest distance value and according to the substitution list of Amino Acid Explorer, Isoleukine is the most common substitution of Leukine. Figure 2 shows the first five most common amino acid substitutions for Leucine according to ‘Common Substitution Tool’ (Figure 2a) and according to the Euclidean distance matrix (Figure 2b). Evidently, the Euclidean distance matrix can predict the most common substitutions of Leucine, but in slightly different order. For example, Common Substitution Tool reports the following order of amino acids: Isoleucine, Methionine, Valine, Phenylalanine and Alanine. According to the Euclidean distance matrix the order is: Isoleucine, Phenylalanine, Valine, Tryptophan and Tyrosine. Methionine is the sixth most common substitution of Leucine, while Alanine is the tenth most common substitution of Leucine according to the Euclidean distance matrix. On the other hand, Tryptophan and Tyrosine are the seventh and the eighth most common substitutions of Leucine according to the Common Substitution Tool. Neglecting the order of the first five substitutions, drawing five amino acids and observing three common is marginally significant (p-value = 0.07, right tail of hypergeometric distribution). Differences between two reports are possibly due to the fact that Euclidean distance matrix is solely based on the structural information of a protein family, while the Common Substitution Tool is based on BLOSUM62 substitution matrix that was created using alignments of multiple protein families.

**Figure 2:**
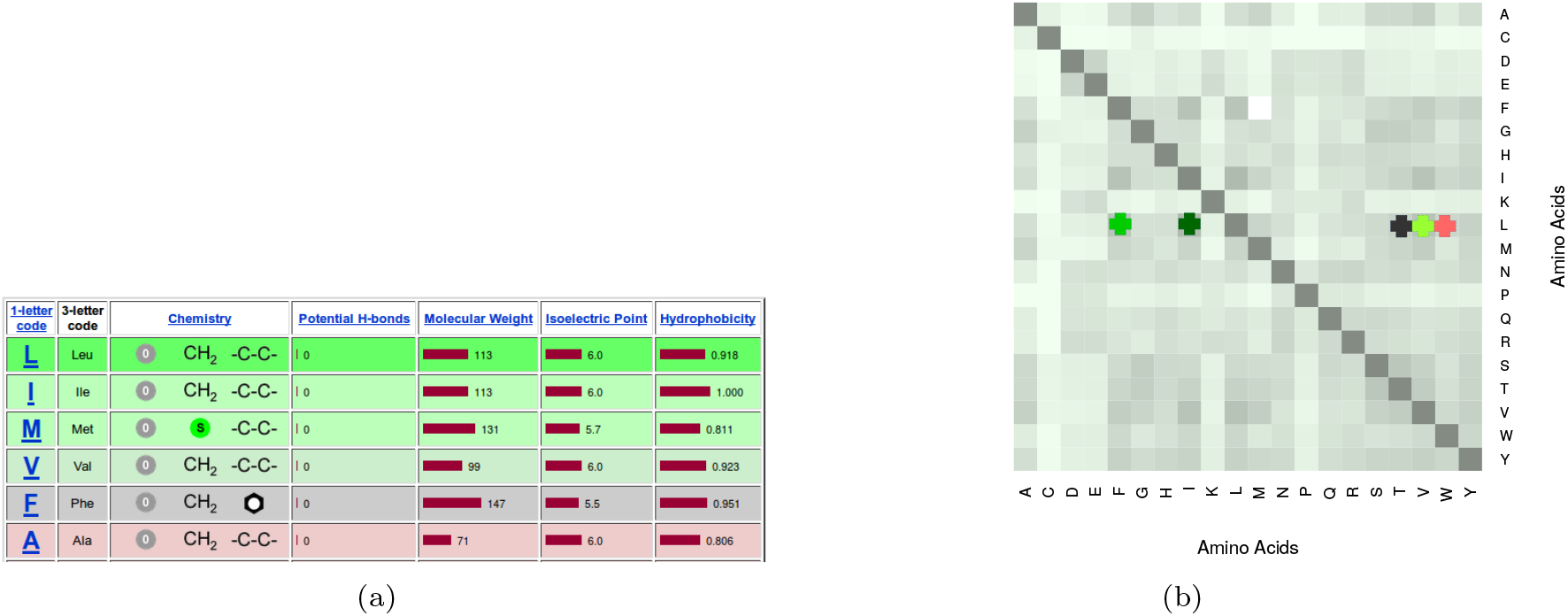
The first five most common substitutions of Leucine according to (a) Common Substitution Tool and (b) Euclidean Distance Matrix. The same color code is used in both figures. Dark green shows the first most common substitution, while the red shows the fifth most common substitution. Methionine and alanine are not found among the first five most common substitutions of Leucine in Euclidean distance matrix, while Tryptophan and Tyrosine are not found among the first most common substitutions in Common Substitution Tool.

According to the previous analyses, we can rely on the structural information of the proteins, which is given by the Euclidean distance matrix, to model amino acid substitutions similarly as using sequence data.

### 3.2 Pairwise comparison of proteins in different species

Using the Euclidean-based distance matrix, we scored the differences of each protein sequence to its human homologue. Results are presented as a heatmap and are clustered hierarchically based on their distance from the human homologue (Figure 3.2). Darker gray tone in Figure 3.2 denotes high similarity between human and other species homologues, whereas lighter tones suggest lower similarity. As expected, proteins from species that are evolutionarily distant from humans are more dissimilar to human homologues. This is especially true for a group of proteins clustered together using the Euclidean distance-based substitution matrix (Figure 3.2 proteins labeled with three dashes). The same analysis was repeated by using the Genetic Code, Manhattan and Pearson distance matrices instead of Euclidean distance matrix. The results are shown in **Supplementary Figures 10a–10c**.

**Figure 3:**
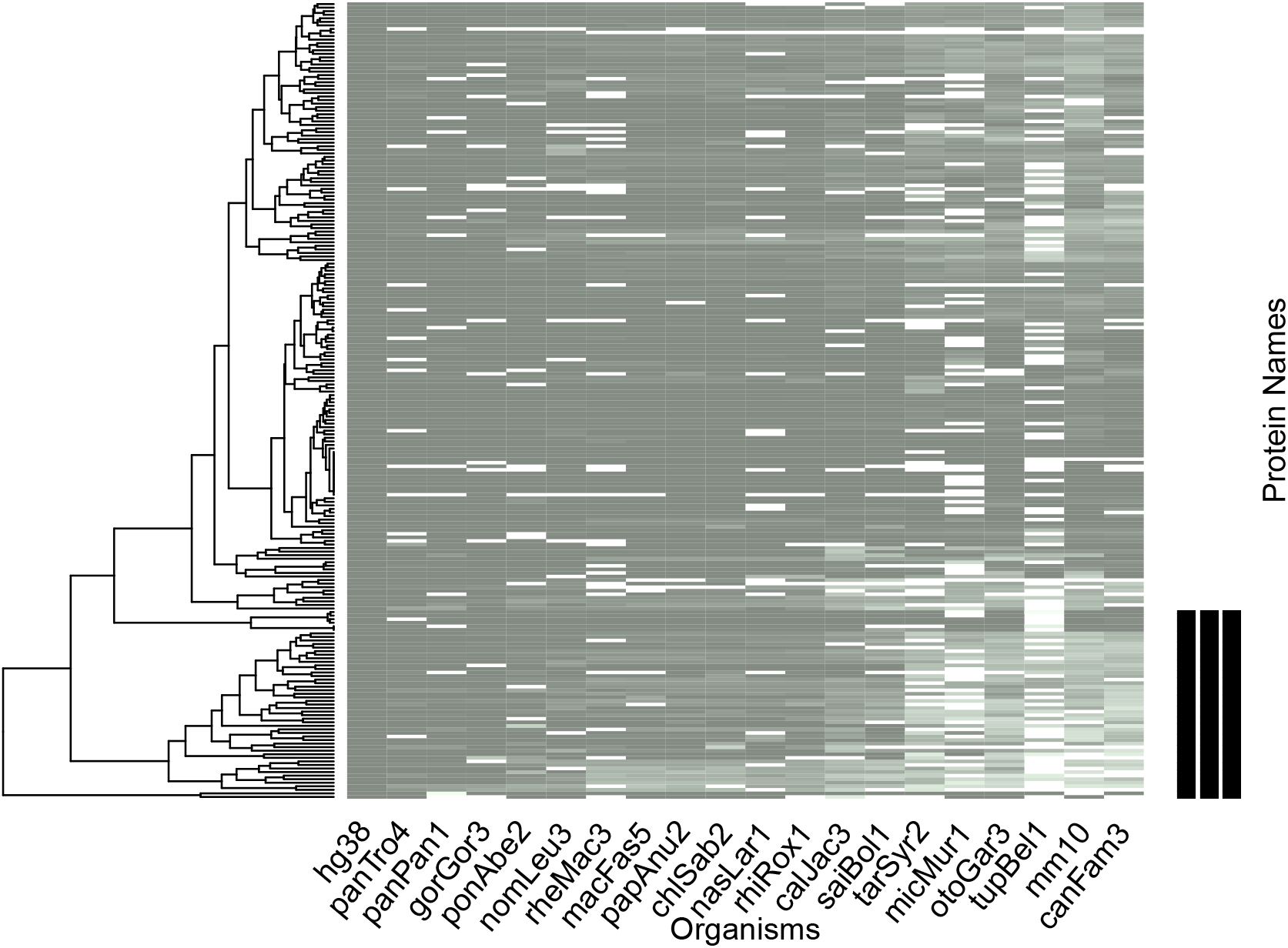
An illustration of the scoring of multiple alignments with the Euclidean distance matrix as a heatmap. Each protein (rows) from human (first column) is compared against the homologous proteins from another species (columns) using pairwise amino acid sequence alignments. We used only the alignment sites with amino acids *A* and *B*, where *A ≠ B* and *A* ≠ − and *B* ≠ −. The total score (a cell in the heatmap) is calculated as the sum of all site scores. Proteins form two clusters based on the hierarchical tree on the left side of the figure. Proteins of the bottom cluster (also denoted with three dashes are further analyzed using Gene Ontology terms.

**Figure 4:**
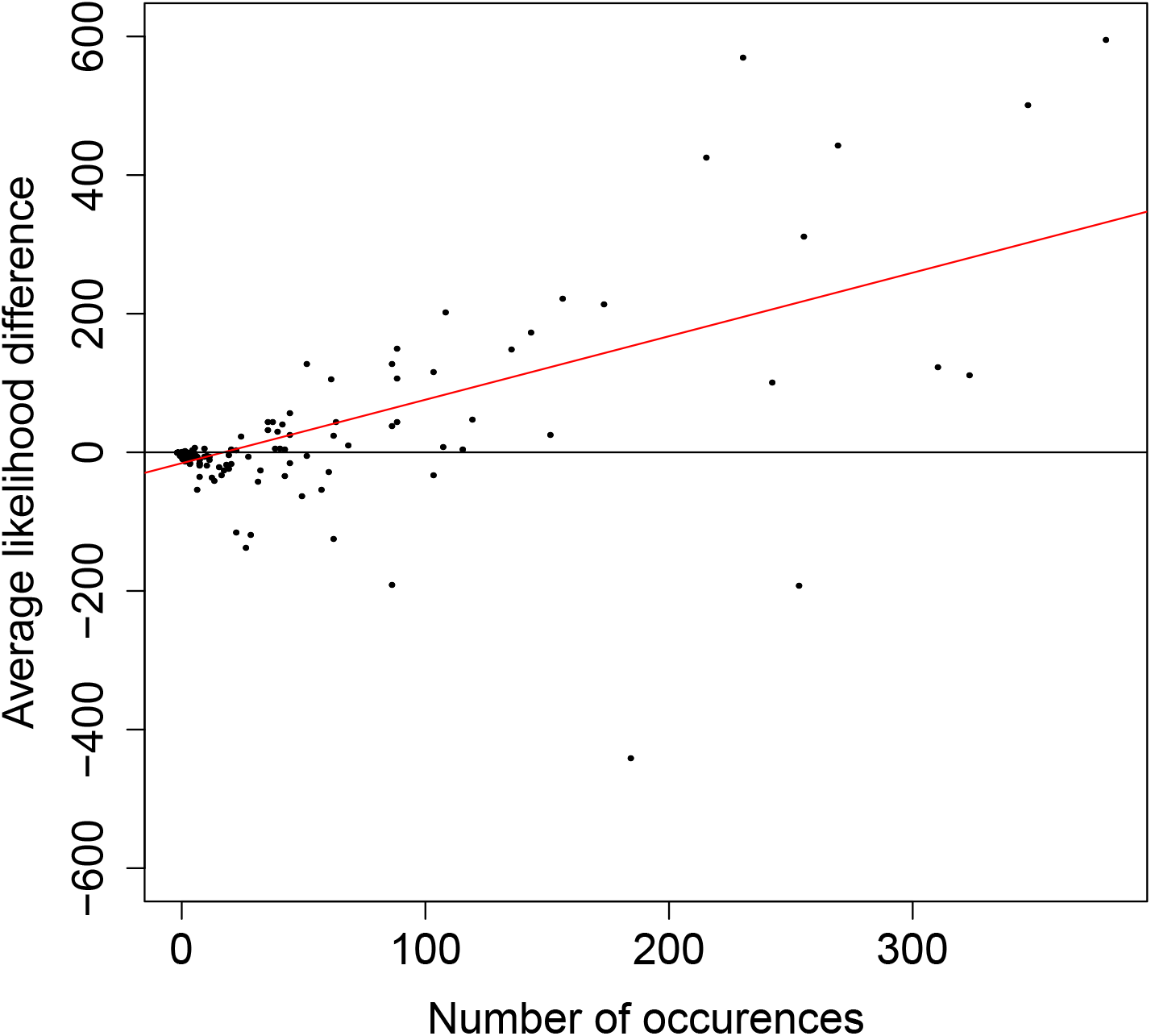
The average likelihood difference between the BLOSUM62-based and the Euclidean-based approaches that have been used to calculate likelihoods. As the figure indicates, for the majority of amino acid pairs the likelihood difference is close to 0, i.e., both the Euclidean-based calculations and the BLOSUM62-based calculations result in similar outcomes. Positive values are fewer than negative values (see text), however the magnitude is much greater for positive values than for negative, indicating that BLOSUM62 results in much greater likelihoods than the Euclidean-based approach. Furthermore, as the occurence frequency of the amino acid increases the difference of the likelihood increases as well, suggesting that BLOSUM62 outperforms the Euclidean-based approach for the amino acid pairs that occur frequently, either within the same protein or in different proteins.

The group of proteins clustered together by the Euclidean-based matrix (marked in the lower part of Figure 3.2) was further scrutinized by Gene Ontology (GO) terms. We used gProfiler (Reimand et al., 2007), to obtain the related GO terms for this set of proteins. These proteins are characterized by a distinct function compared to proteins found in other clusters. On the one hand, the proteins that are located in this cluster play a crucial role in the intestinal absorption of phytosterol and in cholisterol and lipid transportation. On the other hand, the remaining proteins, which are not found in this discrete cluster, are responsible for the homeostatic and transducer/receptor mechanisms of the cells.

Hierarchical clustering of species based on the *nEd* distance matrix orders the species from human as expected, putting Chimpanzees, Gorilla, Orangutan and other primates closer to human, while non-primate mammals (e.g. mouse and dog) are located more distant (Figure 3.2). Thus, using the *nEd* distance matrix, the evolutionary relations between species behave as expected.

### 3.3 Overall and Site Likelihoods

By converting PrInS distance matrices into rate matrices, it was possible to compute the overall likelihood and site likelihoods for every protein alignment. The resulting matrices from the *nEd, nMd* and *nPSm* are named as *nEs, nMs* and *nPSs*, respectively. For most of the proteins, 170 out of 223, the likelihoods computed by BLOSUM62 and PAM120 were better than our neighborhood Euclidean-based substitution matrices. This result is expected since BLOSUM62 and PAM120 substitution matrices were constructed based on alignment files, whereas the presented substitution matrices are alignment-unaware. For the remaining 54 protein alignments, our models and more specifically the Euclidean-based rate matrix resulted in higher likelihoods.

For every protein, we calculated the individual site likelihood for each amino acid site in the multiple sequence alignment using either the BLOSUM62 (or PAM120) or the *nEs* matrix. Results are illustrated in Figure 5) for the protein NCKX1, where the likelihood difference between BLOSUM68 and *nEs* is plotted for every amino acid. Thus, negative values are related to sites where the likelihood is greater for the *nEs* matrix, whereas positive values show sites where BLOSUM62 results in greater likelihoods. For the protein NCKX1, 240 sites are scored with a higher likelihood using the Euclidean substitution matrix derived from PrInS (negative values), whereas 390 sites are scored with a higher likelihood using BLOSUM62 (positive values). The overall likelihood difference for this protein is positive (264.4) indicating that overall BLOSUM62 results in higher likelihood scores. NCKX1 was randomly selected to illustrate the site likelihood differences between the two approaches. It is a critical component of the visual transduction cascade, controlling the calcium concentration of outer segments during light and darkness McKiernan and Friedlander (1999).

**Figure 5:**
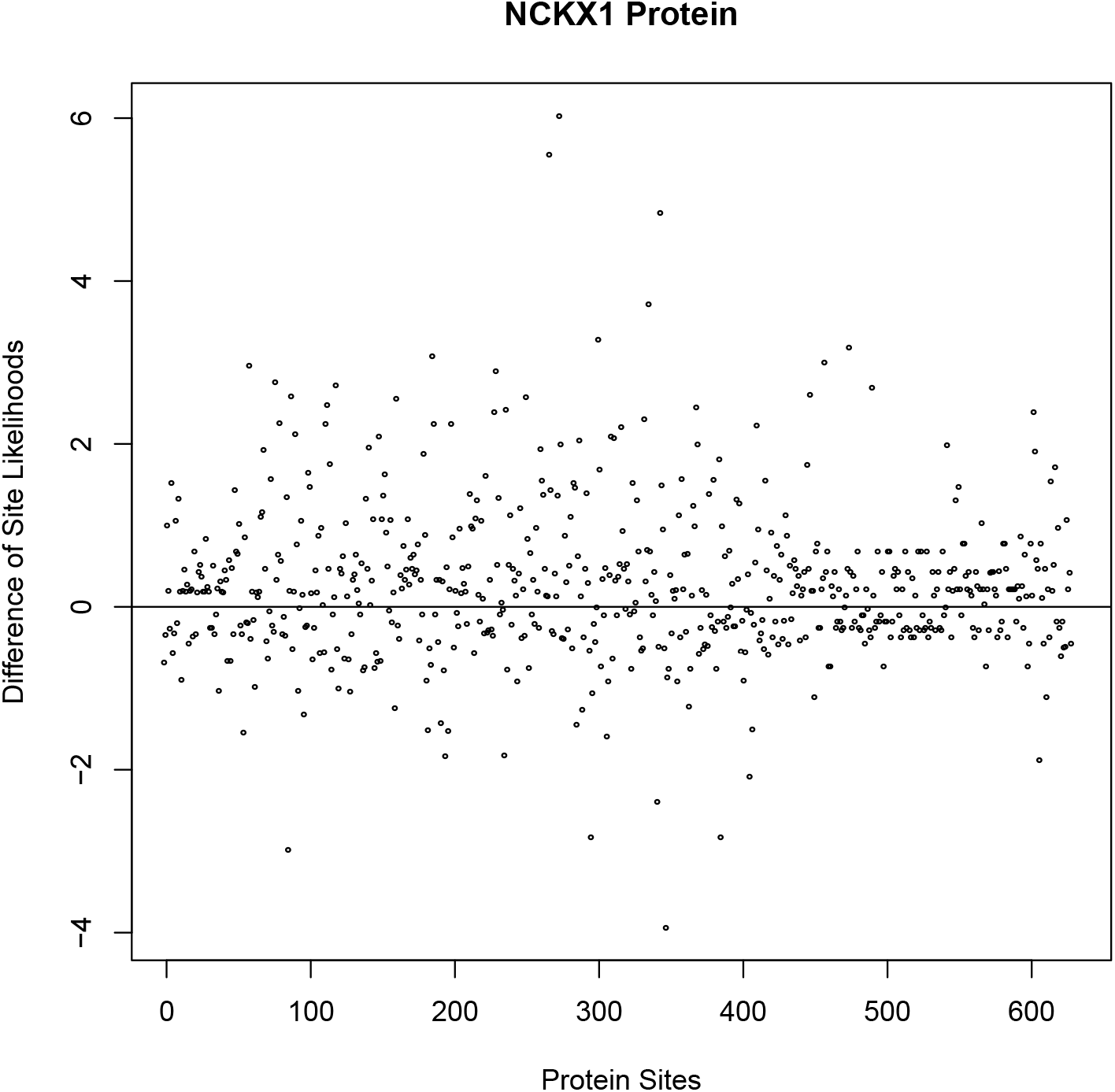
In this graph the differences between the BLOSUM62-based and the Euclidean-based likelihoods for each site of the NCKX1 protein is shown. On the x axis, the protein sites are shown, while on the y axis the differences between the likelihoods are depicted. Positive differences indicate higher likelihoods for the BLOSUM62 rate matrix, whereas negative differences indicate higher likelihoods for the Euclidean rate matrix.

In the **Supplementary Figures 11a–11c** we show the site likelihood differences between the BLO-SUM62 rate matrix and (i) the genetic code rate matrix **11a**, (ii) the Manhattan rate matrix **11b** and (iii) the Pearson rate matrix **11c**.

In calculating overall and site likelihoods, the sequence-based methods, such as BLOSUM62 and PAM120 rate matrices are better than the structure-based methods (Euclidean matrix), however this is expected because BLOSUM62 and PAM120 rate matrices are based on sequence alignments and here are used to score sequence alignments, whereas the structure-based methods we present are alignment-unaware.

### 3.4 Comparison between the average substitution likelihoods for different rate matrices

The 20 amino acids form 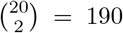 (unordered) pairs. For each of them, we have calculated the average differences between the BLOSUM62-based and the *nEs*-based likelihoods. For the calculations, we considered only the sites that consist of two amino acid states. Since all the parameters of the evolutionary model, but the substitution rate matrix are fixed (phylogenetic tree, equilibrium frequencies), then the matrix that results in the highest average likelihood for a certain amino acid pair describes the preferential model for this amino acid pair. Surprisingly, even though most of the protein alignments are scored with a higher likelihood using BLOSUM62, 111 (out of 190) amino acid pairs are characterized by negative average likelihood difference. This indicates that the *nEs*-based likelihood is greater than the BLOSUM62-based likelihood. The average likelihood is greater for the BLOSUM62 for only 50 pairs, whereas it cannot be assessed for 29 pairs because no occurrences of these 29 pairs were present in the alignment datasets. To scrutinize further this result, we plotted the average likelihood difference as a function of the occurrence frequency of the amino acid pair in the alignment (Figure 3.3). In Figure 3.3, it is apparent that the average likelihood difference is positively correlated with the frequency of occurrences of the amino acid pairs (*r* = 0.621, *r* is the correlation coefficient, CI: (0.55, 0.67), CI denotes the 95% confidence intervals). In other words, the more frequent a substitution is in the alignments, the higher the difference of likelihoods, favoring the BLOSUM62 versus the nEs.

The heatmap in Figure 6 shows all amino acid pairs and the likelihood differences between the BLO-SUM62 and the nEs matrix for all amino acid pairs. Cells denoted by a ‘−’ are characterized by a higher likelihood for the Euclidean-based matrix, whereas a ‘+’ denotes cells with higher BLOSUM62-based likelihood. The darker the color of the cell the higher the difference between the two likelihoods. Cells with an ‘o’ illustrate pairs where no site with this pair of amino acids was found in any of the alignments. The greatest value for a ‘−’ cell is represented for the ‘Arginine-Glycine’ pair. This means that the nEs matrix is a preferential model for the specific amino acid pair. On the other hand, the pair with the highest ‘+’ value is the ‘Isoleucine-Valine’ pair.

**Figure 6:**
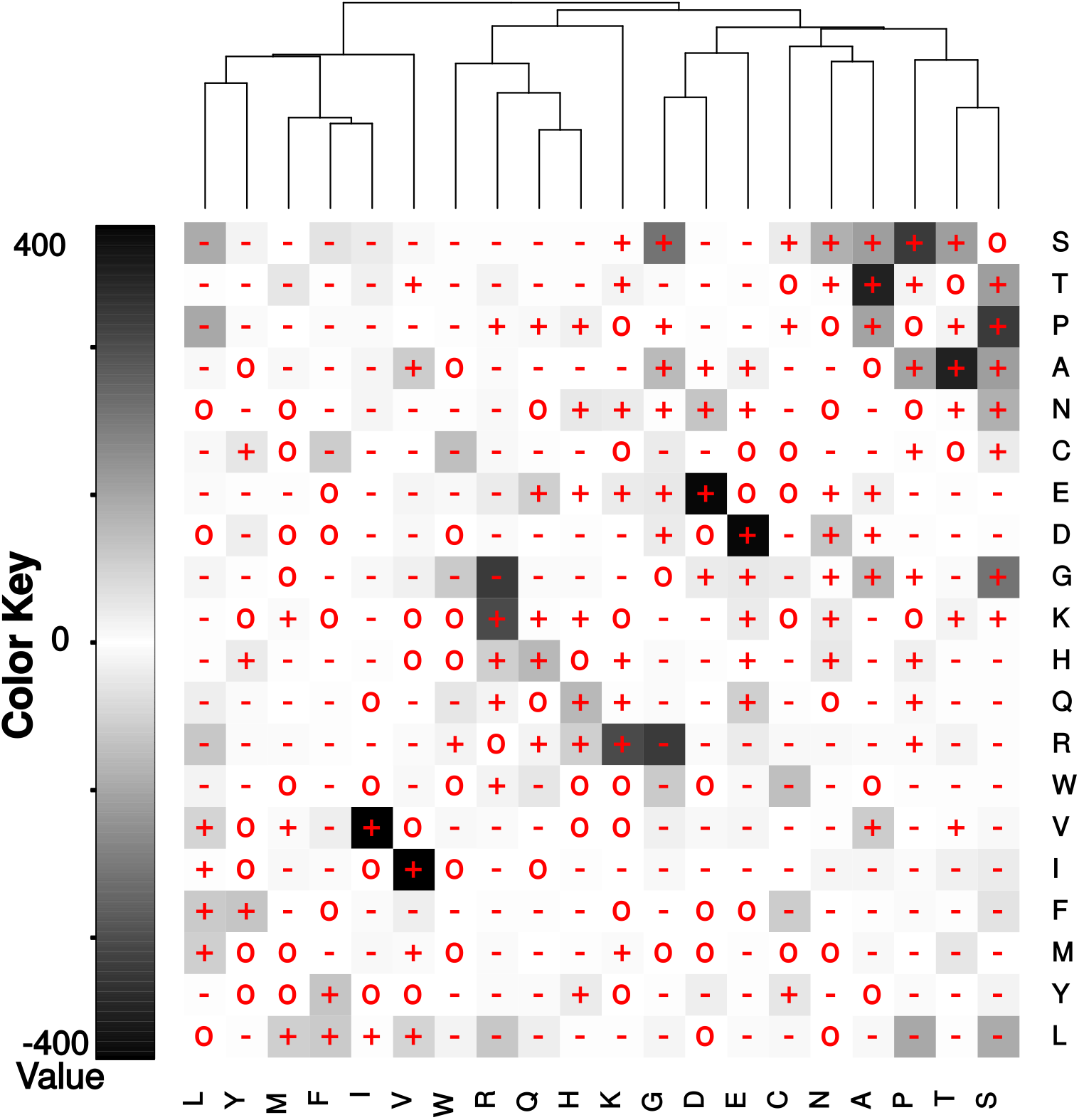
Comparison of likelihoods between BLOSUM62 and *nEs* for all amino acid pairs found in the protein alignments. Boxes with a ‘−’ pinpoint to the specific amino acid pairs where *nEs* calculates a higher site likelihood. Contrarily, a ‘+’ indicates amino acid pairs that the BLOSUM62 approach results in greater likelihood. Finally, ‘o’ cells depict the amino acid pairs that were not found in the multiple alignments, thus no comparison was possible. The darker the tone of the cell the greater the difference in the likelihood between the BLOSUM62 and the *nEs* approach.

In contrast to the overall and site likelihoods, the average substitution likelihoods are calculated better by using structural (Euclidean rate matrix) than sequence (BLOSUM62 and PAM120 matrices) based methods.

### 3.5 Comparison to Grantham’s and Sneath’s distance matrices

Grantham’s distance Grantham (1974) between two amino acids depends on three properties: composition (defined as the atomic weight ratio of noncarbon elements in end groups or rings to carbons in the side chain), polarity and molecular volume. Based on these three properties, distance *D*_*G*[*i,j*]_, between amino acids *i* and *j*, is defined as:

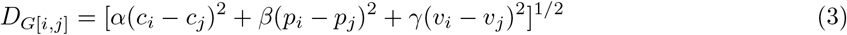

where *c* is the composition, *p* the polarity and *v* the molecular volume. The three components of the distance (*c,p,v*) are not independent. Thus, the constants *α,β,γ* serve as normalizing factors and they can be calculated as a function of *c,v,p* Grantham (1974). Grantham (1974) provides the distances of Equation 3 for all amino acid pairs, thus a distance matrix *D*. Furthermore, Grantham demonstrated that the relative substitution frequency of amino acid pairs and their distances *D*_*G*[*i,j*]_ ‘s are correlated, underlying the physicochemical basis of amino acid substitution.

Similarly to Grantham’s amino acid distances, Sneath’s index Sneath (1966) takes into account 134 categories of activity and structure. The dissimilarity index *D*_*S*[*i,j*]_ is the percentage of the sum of all properties not shared between two amino acids.

The amino acid neighborhood-based distance is correlated with both the Grantham’s and Sneath’s distances Figure 7, highlighting the fact that amino acid neighborhoods capture information related to the physicochemical properties of amino acids and also their relative substitution frequency.

**Figure 7:**
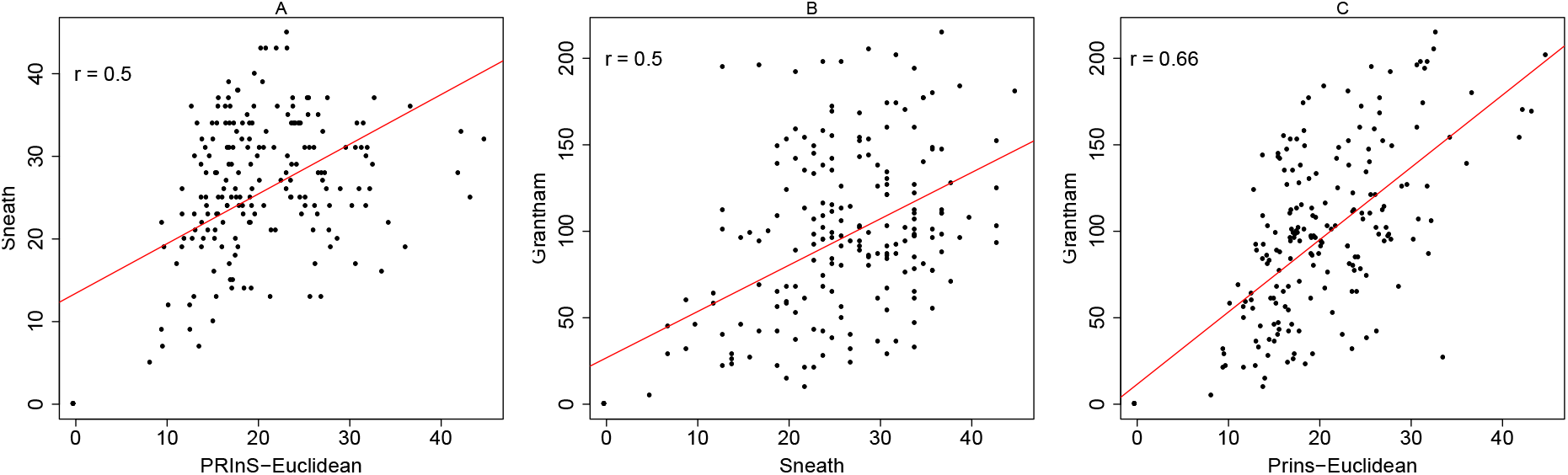
Scatterplots between amino acid distances for different distance methods. (A) Euclidean-based distance versus Sneath’s index (*r* ≈ 0.5). (B) Sneath’s index versus Grantham’s distance (*r* ≈ 0.5). (C) Euclidean-based distance versus Grantham’s distance (*r* ≈ 0.66).

The correlations between the Euclidean and Sneath’s and the Euclidean and Grantham’s distance matrices comprise additional evidence for the ability of protein structural information to predict the amino acid substitutions in a protein family.

## 4 Discussion

### 4.1 Evaluation of PrInS Ability to Predict Amino Acid Substitutions

In this study, we focused on the amino acid substitutions. We investigated if they can be determined from the properties of their neighborhood tertiary structure. We described statistically the amino acid residual neighborhoods using the software PrInS. Then, we converted PrInS output files to amino acid distance matrices and substitution rate matrices and evaluated their ability to model evolutionary changes.

Correlation analysis between the observed substitution frequencies (from multiple sequence alignments) and the distance matrices indicated that residual neighborhoods capture evolutionary information and thus they can be useful in modeling evolution. In other words, substitutions in multiple sequence alignments can be predicted from the amino acid neighborhoods: residuals with similar neighbors can substitute each other, with a higher rate, during evolution. More specifically, from the three distance matrices (Euclidean, Manhattan, and Pearson Squared) that were compared against the substitution matrix, only five amino acids were discordant in at least two comparisons. These amino acids were proline, arginine, serine, tryptophan and tyrosine. From the substitution matrix, only a few substitutions involve tryptophan and tyrosine, whereas there is a multitude of substitutions involving proline, arginine and serine. In contrast, based on distance matrices, in the columns of tryptophan and tyrosine there are small distance values, while in the proline, arginine and serine columns distances are large. This can explain the discordance of these amino acids in these two matrices, because a large number of substitutions is associated with small distance values for an amino acid.

In addition, the Common Substitution Tool of NCBI Amino Acid Explorer (Bulka et al., 2006) enhances our main result: Amino acid pairs with the lower distance values in the distance matrices, especially using Euclidean distances, are found to substitute each other more frequently. Amino Acid Explorer returned a list of the amino acids from the most common to the the less common substitution for an amino acid we put as input. There are some differences in the order of the amino acids between the Common Substitution Tool and the distance matrices (e.g Euclidean). These differences are possibly based on the fact that the generation of the Euclidean distance matrix is based solely on the structural information of a protein family (*α*-helical membrane proteins), while the Common Substitution Tool is based on BLOSUM62 substitution matrix that was created from alignments from multiple protein families. This possibly means that some specific amino acid substitutions are observed more frequently in the *α*-helical membrane proteins, which are not found so often in the other protein families. These substitutions may play a crucial role in the distinct function of the *α*-helical membrane proteins.

### 4.2 Multiple Alignment Scoring and Clustering of Proteins

After the evaluation of the ability of PrInS to predict amino acid substitutions, the scoring of multiple alignments with the matrices generated by our methodology was followed. The matrices that were used in the scoring of multiple alignments were distance-based, genetic-code-based and model substitutions matrices (BLOSUM62 and PAM120) Henikoff and Henikoff (1992), Dayhoff et al. (1978). This was the second step in our analysis. Generally, most of the proteins did not significantly differ from the human homologues, which are found in the first column of all the heatmaps. Obviously, homologous proteins in evolutionary distant species from human differ more than the homologues in more related species. Based on the substitution rate matrices, we calculated the likelihood for every protein pairwise alignment (human *vs* non-human species). Using the Euclidean-based matrix, a cluster of proteins was appeared (hierarchical clustering) and its functions were studied using gProfiler Reimand et al. (2007). The proteins in the discrete protein cluster play a crucial role in the intestinal absorption of phytosterol and in cholesterol and lipid transportation, while the proteins of the other clusters are responsible for the homeostatic and transducer/receptor mechanisms of the cells.

## 5 Conclusion

The main result of this project is that, by using only protein structural data, we are able to predict the amino acid substitutions in a protein family. In many cases these results are similar with the results by using only sequence data. Finally, based on these outcomes we can support our initial hypothesis that the amino acids with the same amino acid neighbors can substitute each other during evolution.

## 6 Supplementary Figures

**Figure 8:**
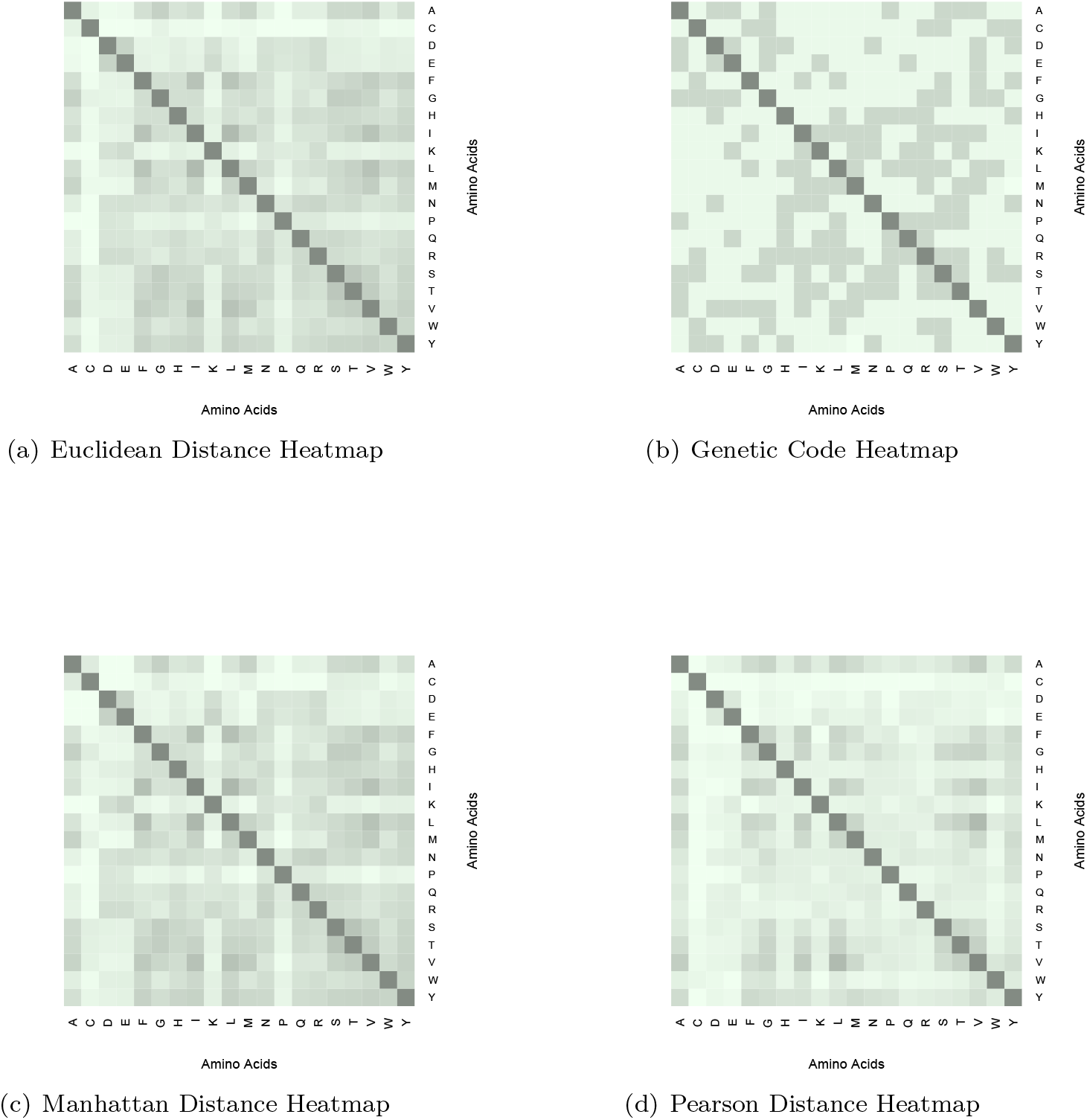
Heatmaps that depict (a) the Euclidean Distance Matrix, (b) the Genetic Code Matrix, (c) the Manhattan Distance Matrix and (d) the Pearson Distance Matrix. (a) In this figure, such as in figures 6.1c and 6.1d (Distance Matrices) the lower distance values are depicted with dark grey, while the larger distance values with the with white. Again the lower distance values in amino acid pairs denote ”behavioural” similarity. (b) In the genetic code heatmap, the fewer base changes are depicted with the grey colour, while the many base changes with white. Finally, this heatmap shows how many base changes have to be occurred in order to change an amino acid into another. (c-d) As it is obvious, there are fewer behavioural amino acid similarities in the Pearson heatmap than in Manhattan distance heatmap.

**Figure 9:**
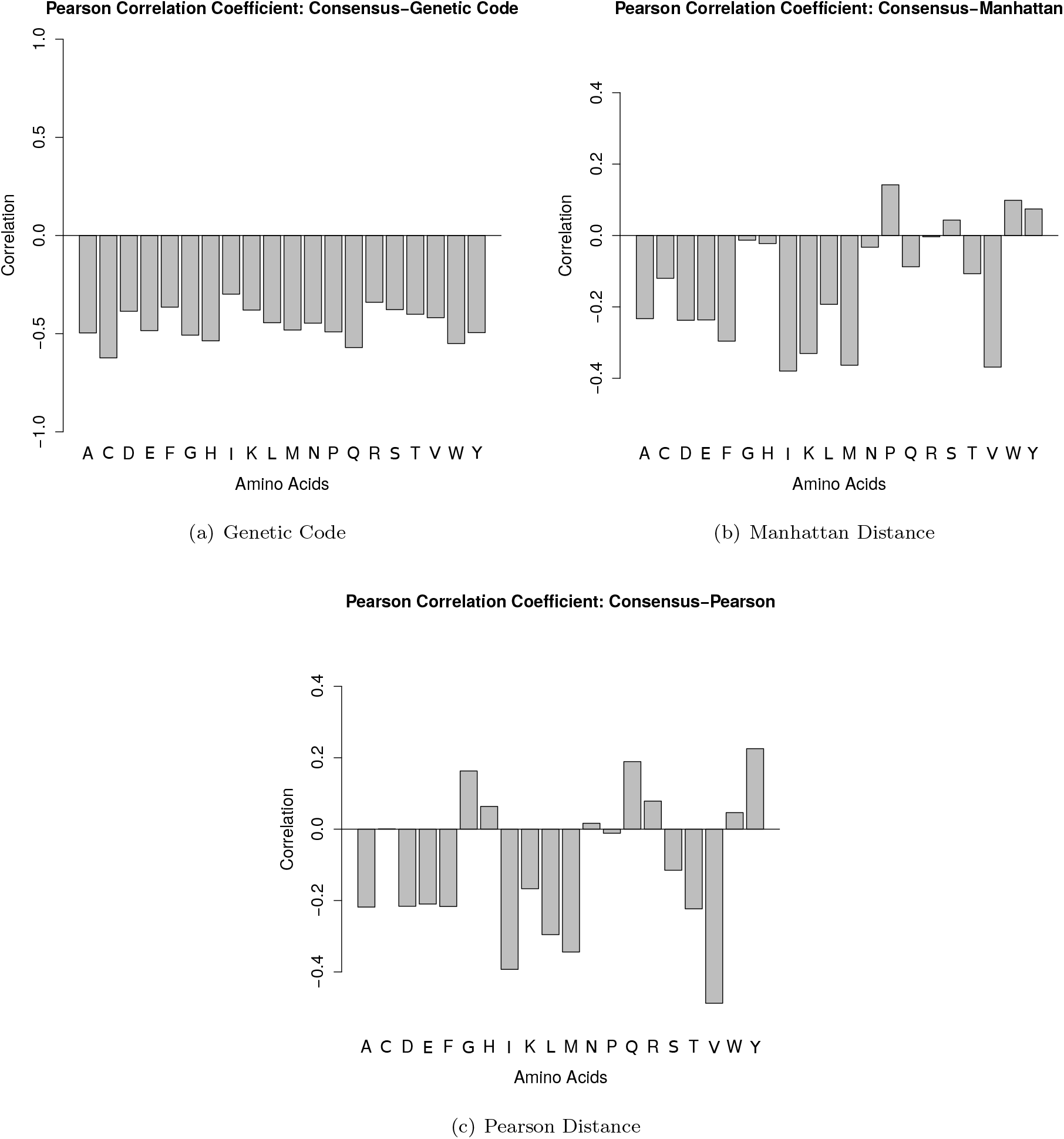
Pearson Correlation Coefficient between the consensus and (a) Genetic Code Matrix, (b) Manhattan Distance Matrix and (c) Pearson Distance Matrix. The correlation is represented with the negative values, while the disassociation with positive values, due to the different nature of consensus and genetic code/distance matrices.

**Figure 10:**
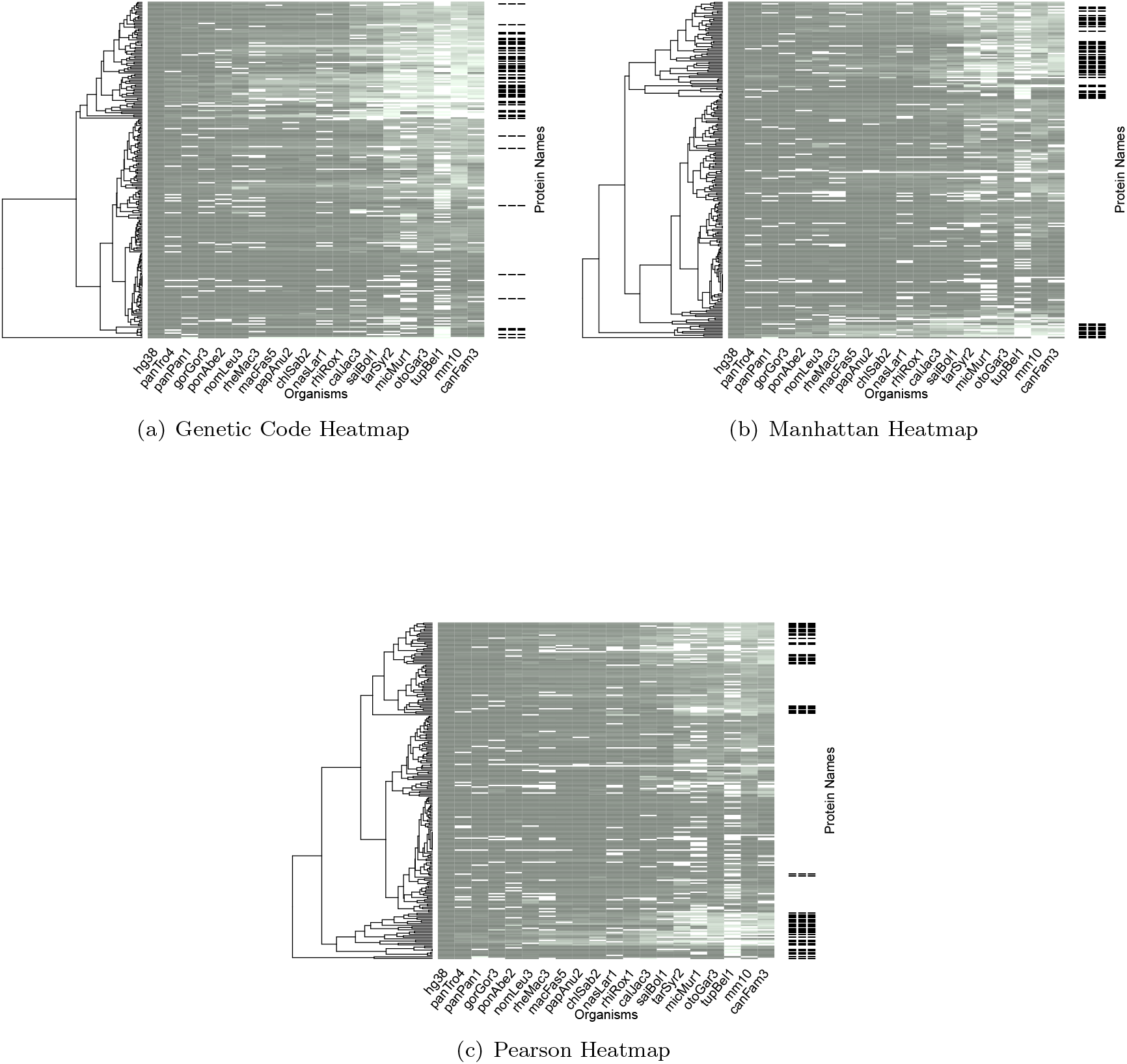
Heatmaps that depict the scoring of multiple alignments with the (a) Genetic Code Matrix, (b) Manhattan Distance Matrix and (c) Pearson Distance Matrix. In this figure there are also shown the hierarchical clustering of proteins and the species, whose proteins were scored. Human is found in the left end of the heatmaps. The other species were put in evolutionary order. Finally, the proteins that were found on the protein cluster in Euclidean heatmap (Figure 3.2a in the main part of the manuscript) are represented in these heatmaps with three dashes. In these heatmaps the clustering of the proteins that were found in Euclidean heatmap is more obvious than in BLOSUM62 and PAM120 heatmaps (Figure 3.2b and 3.2c in the main part of the manuscript).

**Figure 11:**
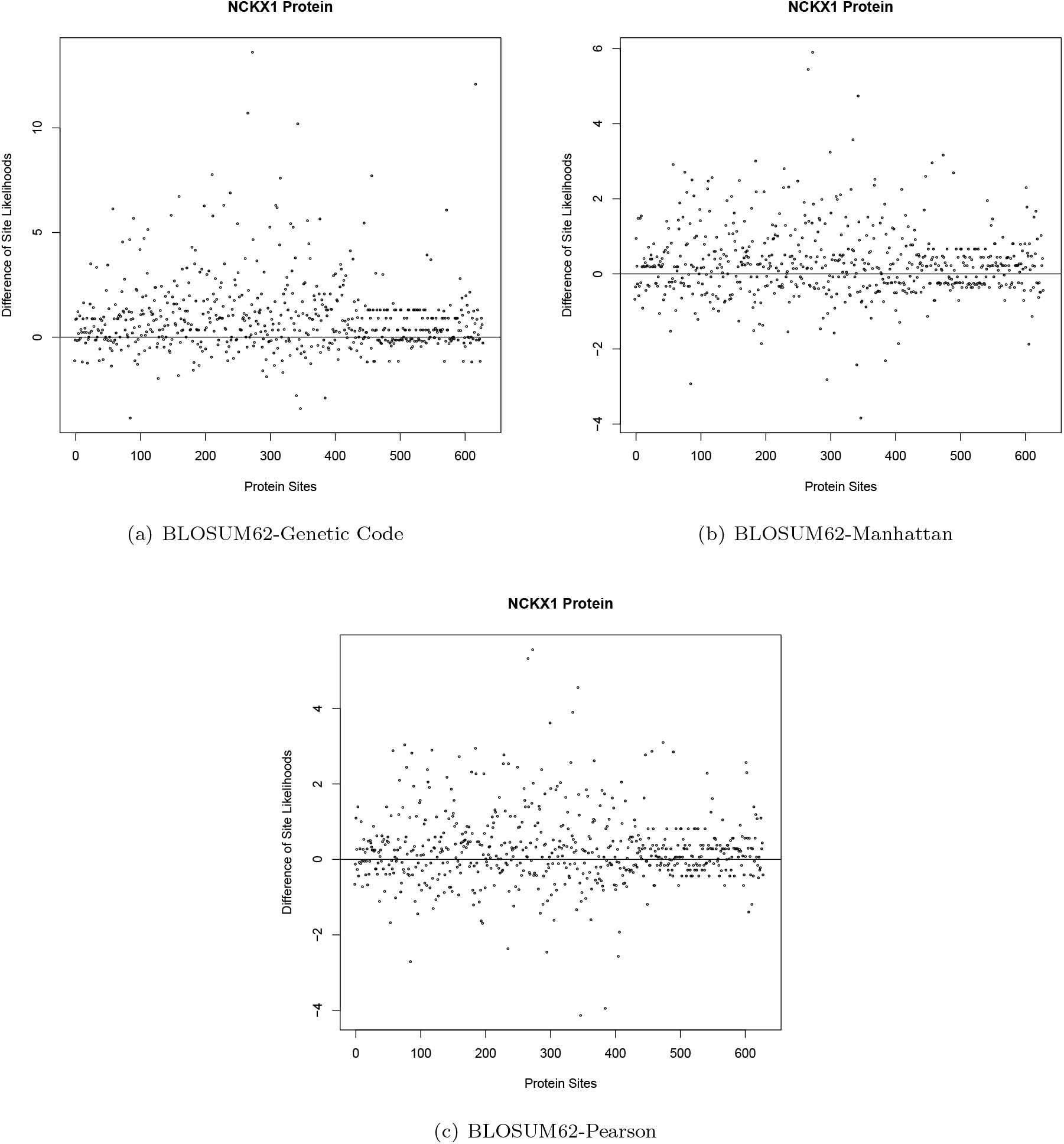
In these graphs the differences between the BLOSUM62 and the (a) Genetic, (b) Manhattan and the (c) Pearson Site Likelihoods for the NCKX1 protein are shown. On the x axis, the protein sites are shown, while on the y axis the differences between the likelihoods are depicted. On the one hand, positive differences indicate that the BLOSUM62 rate matrix is more reliable in the calculation of these specific site likelihoods. On the other hand, negative differences indicate that the (b) Genetic Code, (c) Manhattan and (d) Pearson rate matrices more reliable in the calculation of these specific site likelihoods.

